# Peptidoform analysis of IP-MS data allows detection of differentially present bait proteoforms

**DOI:** 10.1101/2024.01.23.576810

**Authors:** Savvas Kourtis, Damiano Cianferoni, Luis Serrano, Sara Sdelci

## Abstract

While it is recognised that protein functions are determined by their proteoform state, such as mutations and post-translational modifications, methods to determine their differential abundance between conditions are limited. Here, we present a novel workflow for classical immunoprecipitation coupled to mass spectrometry (IP-MS) data that focuses on identifying differential peptidoforms of the bait protein between conditions, providing additional information about protein function.

## Main

Proteins are widely regarded as the functional units of cells, facilitating functions such as transcription, translation, metabolism and signal transduction. The specific functions that each protein performs are determined by its structure, which is in turn encoded by its amino acid sequence. Mutations to this sequence, as well as post-translational modifications can give rise to proteoforms that expand the repertoire of functions that a protein can perform. Such changes in proteoform function include phosphorylation-based activation, localisation, mutation-mediated oncogenicity among others^1^.

Recent efforts have attempted a genome-wide cataloging of proteoforms in specific conditions such as blood cell types^2^, which opens the possibility of determining differential proteoform presence. Top-down proteomics has been demonstrated to detect differentially present proteoforms associated with liver transplant rejection^3^. However, despite advances in the field, top-down is still not widely adopted by the community, creating the need for a bottom-up differential proteoform approach.

Here we present a workflow for the detection of differentially present peptidoforms^4^ from classical IP-MS experiments, which traditionally aim to detect the interactome of a bait protein^5^. While classical IP-MS focuses on the differential peptide spectrum matches (PSMs) of ‘prey’ proteins to identify interactors, we propose that the differential PSMs of the ‘bait’ protein could provide differential peptidoform information (Fig 1A). Our workflow is based on the premise that the bait peptidoforms are enriched by the antibody-based capture of the bait by the protocol, allowing their reproducible detection through MSFragger Open search^6,7^ and differential analysis using SAINTexpress^8^. For the very first time, our approach eliminates the need for PTM-specific enriched samples and allows simultaneous capture of modified bait peptides (Fig 1A).

**Fig 1:**
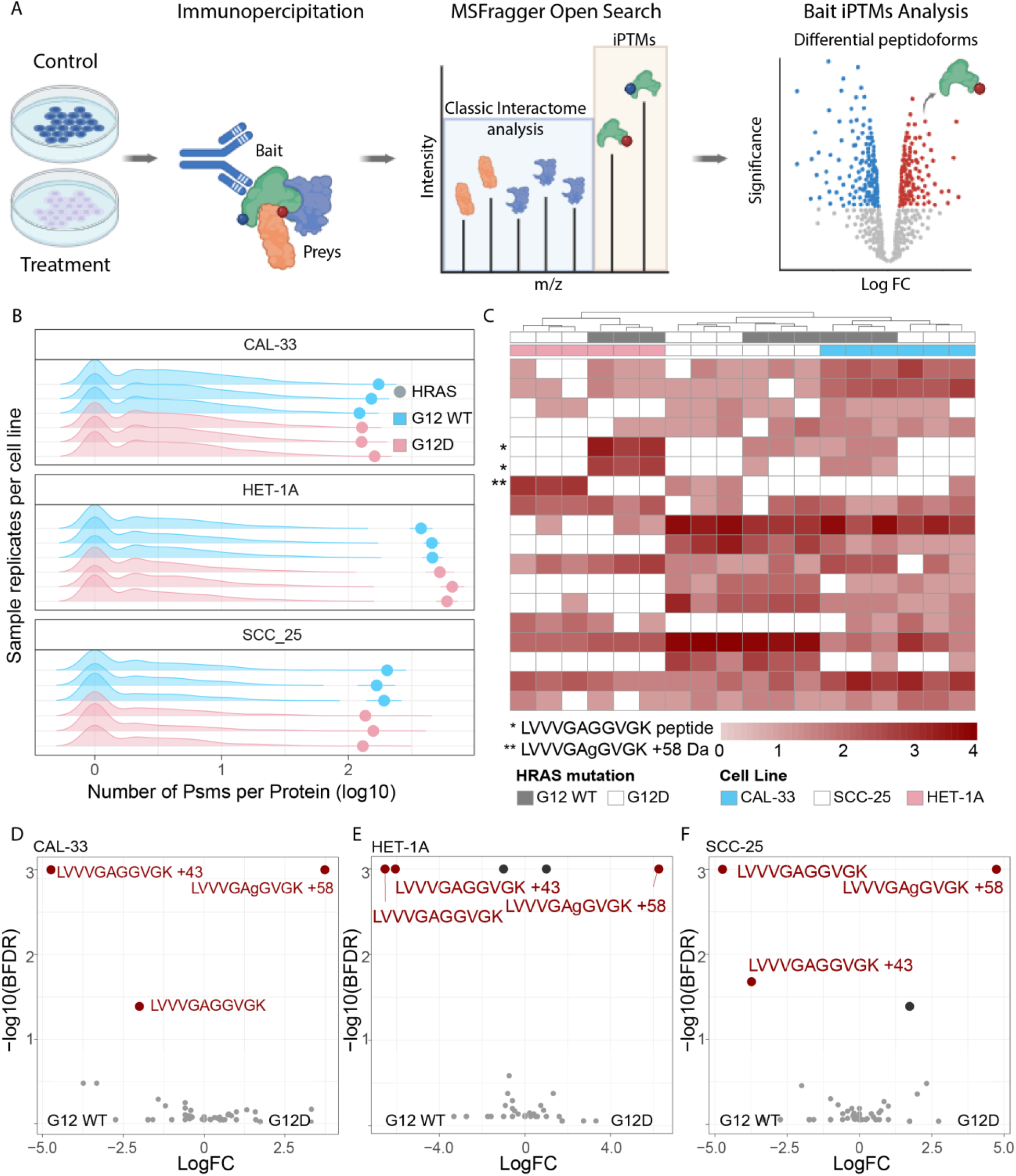
a. iPTMs workflow for analysis of IP-MS data, using MSFragger in Open search with SAINTexpress downstream to identify differentially present peptidoforms between conditions. b. Pre-normalisation PSM counts for prey (density plots) and bait (dots) proteins for paired HRAS WT (blue) and G12D (salmon) samples across cell lines, for each replicate IP-MS experiment. c. Clustering of post-normalised PSM counts for highly abundant peptidoforms belonging to bait HRAS protein. Unsupervised clustering performed with default hclust parameters with peptidoforms not detected in specific samples assigned a zero value following log transformation. d. D-F differential bait peptidoforms detected between WT and G12D HRAS in CAL-33 (D), HET-1A (E), SCC-25 (F) cell lines according to the SAINTexpress, defined as BFDR< 0.05. Only bait peptidoforms were included in the SAINTexpress analysis.

By re-analysing pulldowns of exogenously expressed HRAS wild-type (WT) and G12D mutant constructs in 3 cell lines (PXD019469), we showed that the HRAS bait proteins are highly enriched in these samples and are detected by numerous PSMs, achieving 98% bait protein coverage (Fig 1B, S1A). Many of these peptides were only captured in a modified state, and thus would not have been detected using a conventional closed search (Fig S1B), therefore decreasing the total protein coverage. Furthermore, we showed that the bait PSMs correctly clustered samples both by cell line and HRAS mutation status, highlighting that our methodology captures biologically relevant HRAS states between samples (Fig 1C). Among these, the LVVVGAGGVGK peptide is detected in its unmodified, carbamylated (+43), and G12D mutated (+58) forms, thanks to the open search (Fig 1B). The presence of the LVVVGAGGVGK +58 G12D peptide, although not detected in the original study, validates the researchers’ experimental design and results^9^. This demonstrates that IP-MS data coupled with open search, provides additional information about the bait protein that would otherwise be missed. To systematically quantify these changes in the bait protein, we apply the SAINTexpress method (Teo et al., 2014) for each cell line separately. We identified the peptides LVVVGAGGVGK (unmodified) and LVVVGAGGVGK (+58.00 Da G->D substitution) as differentially present between HRAS WT and G12D cell lines (Fig 1D-F). Additionally, MSFragger localised the +58 modification on the G12 position, which is indicated in lower case in the peptide, providing further evidence and interpretability for this modification (Fig 1D-F). Finally, the position of differentially modified peptides could be indicative of an isoform switch, and as expected, when mapping the differential peptides between HRAS WT and HRAS G12D in CAL-33, we observe their strong localisation at the N-terminal of the protein with little differences downstream of the G12 position (Fig S1C). Such positive controls highlight the power and reproducibility of our approach.

The recent discovery and application of the KRAS G12C specific inhibitors has been particularly promising. However, emerging resistance to AMG-510^10^ has already been already reported^11^ and has become a new area of cancer research. Therefore, we investigated the differential peptidoforms of exogenously expressed KRAS WT and G12C upon AMG-510 treatment (PXD043536)^12^. We checked the quality of the IP-MS data by verifying that KRAS was highly enriched in the samples (Fig 2A) and a that high protein coverage was achieved (Fig S2A-B). Interestingly, the addition of AMG-510 appeared to decrease KRAS PSMs, which in the original study was attributed to the inhibitor preventing KRAS trypsinization (Fig 2A). WT KRAS was not strongly affected by the treatment, consistent with the specificity of AMG-510 for G12C mutated KRAS (supplementary table 1). KRAS G12C bait PSMs correctly clustered treated and control samples demonstrating that the treatment with the AMG-510 induced changes to the KRAS G12C proteoform state. Numerous modified peptides were significantly different for KRAS G12C after the AMG-510 treatment (Fig 2B), confirming a change in KRAS G12C proteoform state. Among others, the C118 position (Figure 2C), which is known to be oxidized to allow the release of GDP and contribute to oncogenicity^13,14^, appeared to be affected by the AMG-510 treatment. In particular, C118-modified containing peptides were identified as differentially presented in the samples, being modified to cysteic acid (+47.98 Da) and to a sulfinic acid (+32.00 Da) in presence of AMG-510. The respective -9 and -25 delta mass were most probably due to the absence of carbamidomethylation on these cysteine residues (+57.02), a chemical derivative which arises when samples are treated with iodoacetamide^15^.

**Fig 2:**
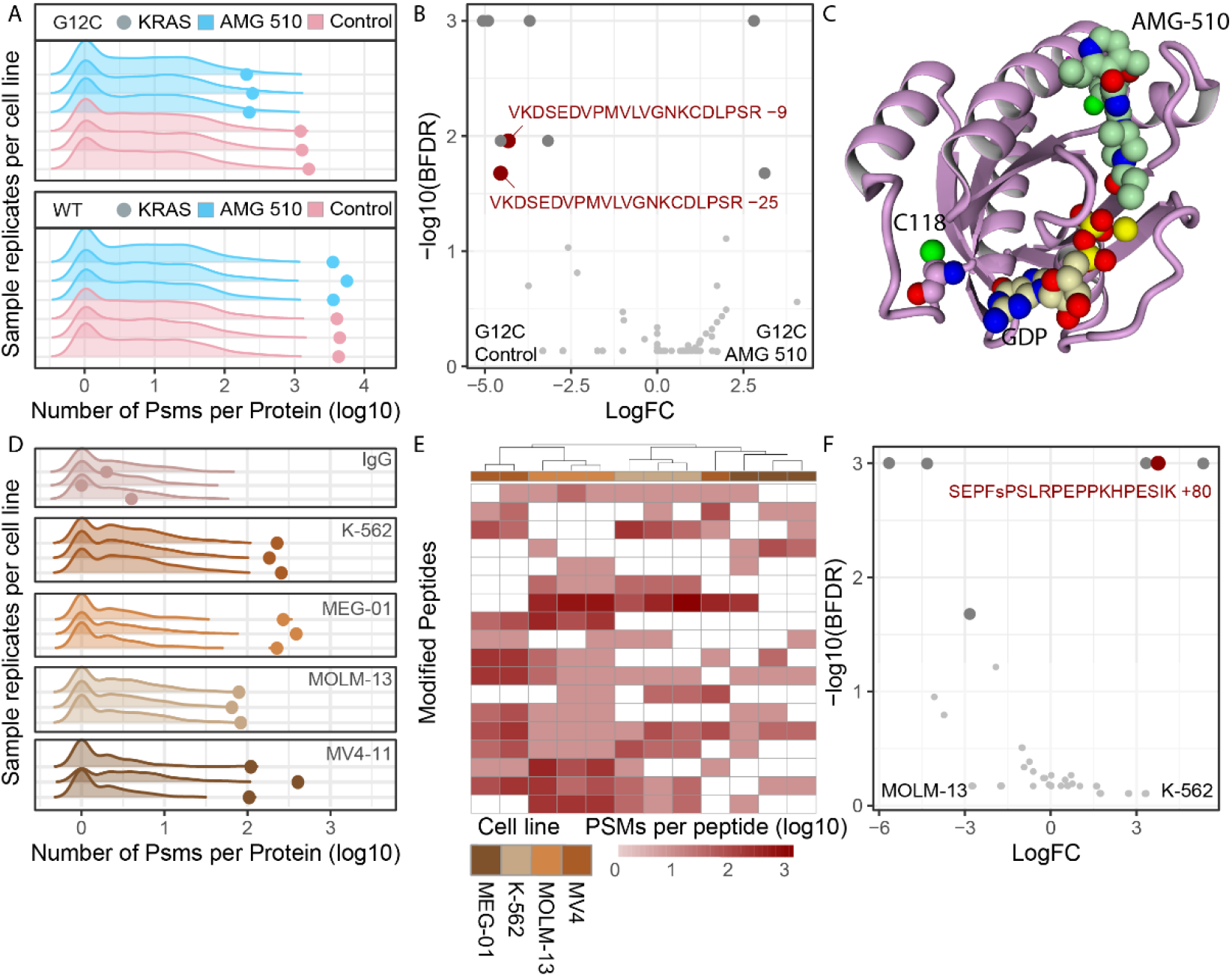
a. Pre-normalisation PSM counts for prey (density plots) and bait (dots) proteins for AMG-510 treated (blue) and untreated (salmon) samples for KRAS WT and G12C mutant cell lines, for each replicate IP-MS experiment. b. Differential bait peptidoforms detected between AMG-510 and untreated KRAS G12C samples according to the SAINTexpress, defined as BFDR< 0.05. Only bait peptidoforms were included in the SAINTexpress analysis. c. Structural model of C118 position relative to AMG-510 and GDP binding site in KRAS G12C protein. d. Pre-normalisation PSM counts for prey (density plots) and bait (dots) proteins for IgG control (top) and endogenously enriched BRD4 across cell lines, for each replicate IP-MS experiment. e. Clustering of post-normalised PSM counts for highly abundant peptidoforms belonging to bait BRD4 protein, excluding the IgG control samples. Unsupervised clustering performed with default hclust parameters with peptidoforms not detected in specific samples assigned a zero value following log transformation. f. Differential bait BRD4 peptidoforms detected between K-562 and MOM-13 samples according to the SAINTexpress, defined as BFDR< 0.05. Only bait peptidoforms were included in the SAINTexpress analysis.

Finally, although exogenously expressed proteins increase yield in IP-MS experiments, they can also increase false positives by changing the physiological state of the bait protein^16,17^. To validate our approach in an endogenous setting, we explored endogenous BRD4 pulldowns in four different leukemia cell lines (PXD012715)^18^. BRD4 was enriched in all conditions except the IgG control (Fig 2D). The different levels of enrichment suggested differential BRD4 expression across cell lines. Our protocol mitigated this effect by normalizing all bait PSMs per sample to make the comparison equitable. Despite the lower protein coverage (Fig S2D), BRD4 bait PSMs correctly clustered 15 out of 16 samples, demonstrating the different proteoform states between cell lines (Fig 2E). Focusing on the K-562 and MOLM-13 cell lines, we identified the differential expression of a phosphorylated peptide (Fig 2F), which underlines the power of our approach in detecting PTMs of endogenously expressed proteins. Interestingly, the majority of BRD4 differential peptides seemed to map to the C-terminus of the protein with few peptides detected in the N-terminus (Fig S2E). Lastly, while it is possible to detect differentially present unmodified peptides (Fig S2F), in the absence of their modified counterpart, these peptides are harder to interpret, as it is unclear whether they represent an isoform switch or perhaps a change in cleavage site.

In conclusion, our methodology allows for the very first time the capturing of differentially modified peptides by analysing antibody capture protocols. Obtaining paired interactome and proteoform changes from the same samples, opens up the possibility of integrating such information with protein-protein interaction (PPIs) interface studies to determine whether the peptidoform changes could interfere with PPIs. However, this approach is not applicable to interactome approaches that directly enrich prey proteins, such as Bio-ID^16,19^.

While our methodology complements classical interactome analysis and can be used to retrospectively analyse publicly available datasets, we hypothesize that protocols can be optimised by increasing the stringency of sample washes to shift the balance of PSMs retrieved towards the bait, enhancing the power of our methodology. Similarly, improved protocols that limit contaminants in IP-MS samples such as anti-fouling agents^20^ would improve proteoform detection. Although not often recognised even in classical IP-MS, the degree of bait enrichment in each sample strongly affects downstream analysis, as in the demonstrated case of BRD4 expression where MV4-11 had low reproducibility in bait enrichment, thus decreasing interactor and peptidoform enrichment in some samples (Fig 2D).

Similar to other trypsinisation-based bottom-up methodologies, our approach cannot directly determine whether the modified peptides co-occur on the same protein molecule. Finally, since bait capture is based on antibody binding to the bait, any PTMs, compounds or interactions that interfere with this binding will prevent these proteoforms from being captured, and a polyclonal capture could be considered to mitigate this.

## Methods

### Data Processing

Raw files were downloaded from the PRIDE proteomics repository^21^, for the HRAS (PXD019469), KRAS (PXD043536) and BRD4 (PXD012715) respectively. These data were analysed using MSFragger (FragPipe 19.2-build 11, MSfragger 3.8, philosopher 5.0.0) open search default settings^6^ with additional information about the mass shifts derived from PTM-Sheppard^22^. Oxidation of methionine was included a variable modification, and cysteine carbamidomethylation was included as a fixed modification according to the original reagents and searches performed in the respective original studies.

### Differential peptide analysis

PSMs mapping to the bait protein were counted per sample per modification state with each peptide being assigned a modification mass of its reported delta mass, rounded to the closest decimal point. The MSFragger mass shift localisation and PTM-Sheppard annotation was used to explore the identity of the mass shift, with larger combinatorial modifications being much harder to characterise. To ensure enrichment differences between samples did not cause bias in the differential analysis, each sample’s total bait PSM count was normalised to the total counts of the sample with the least bait PSMs within each comparison to be made. These normalised counts were the input to SAINTexpress^8^, for differential peptide analysis. Code and file availability at github.com/SdelciLab/iPTMs.

### KRAS AMG-510 structure

Figure was obtained using yasara^23^ visualization software. The model (6oim) was obtained from the Protein Data Bank^24^ and Serine 118 was mutated to Cysteine using FoldX 30874800 BuildModel command^25^. The resulting DDG of 0.81 kcal/mol demonstrates the suitability of the used model to host the wildtype Cysteine residue. The protein backbone represented as pink ribbon, Cysteine 118 is represented as atom spheres (pink carbons), as well as AMX (green carbons) and GDP (beige carbons).

---

**Fig S1:**
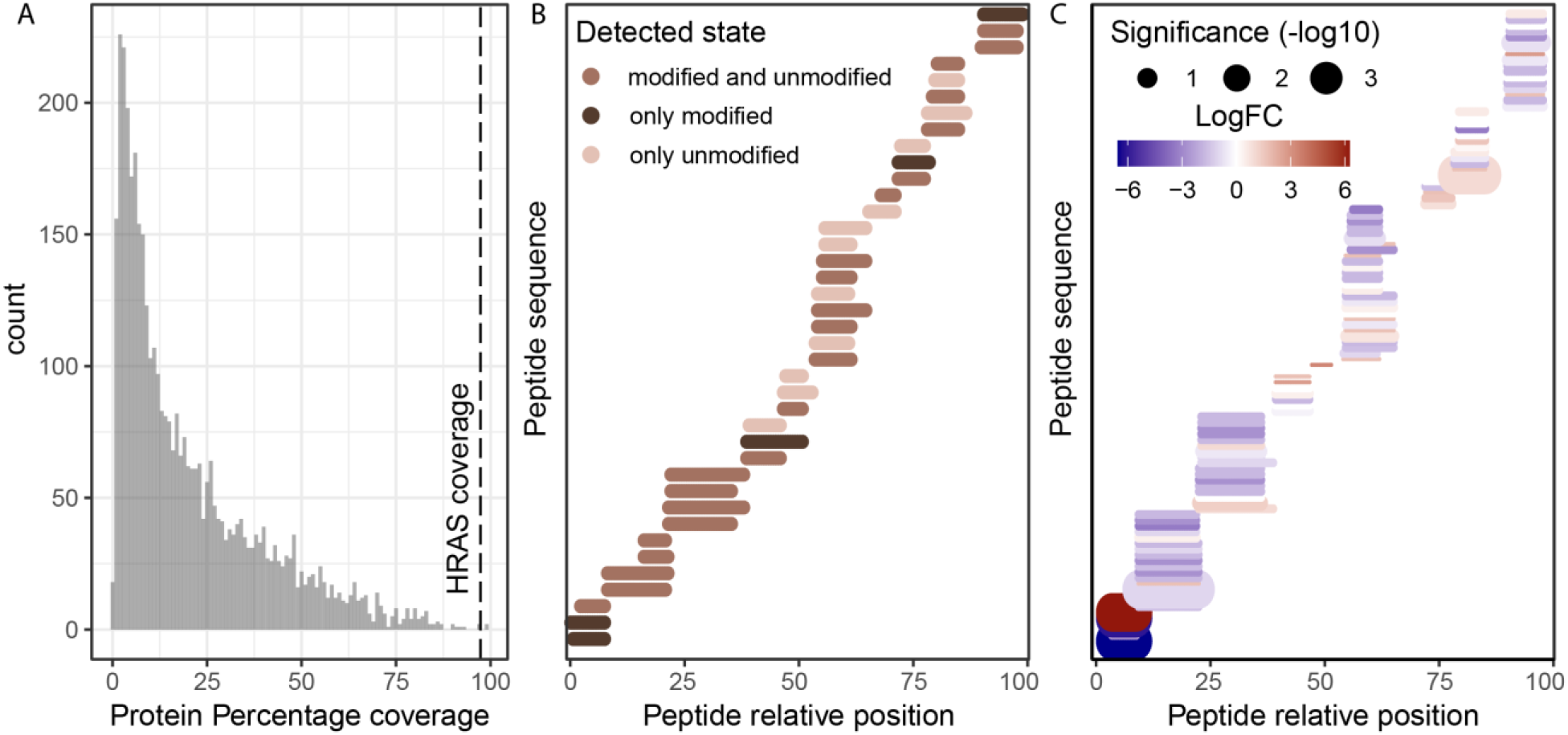
a. Percentage protein coverage for all preys and baits calculated as the percentage of amino acids detected per gene according to the longest isoform, across all HRAS IP-MS samples and all peptidoforms. Dotted line highlight the bait (HRAS) protein coverage. b. Relative position of all detected HRAS bait peptides, as well as whether they were detected in unmodified, modified or in multiple delta masses. Peptides with isotopic error delta masses were considered unmodified for this representation. c. Relative positions of CAL–33 bait peptides with peptides with statistical significance and fold change encoded in size and color respectively.

**Fig S2:**
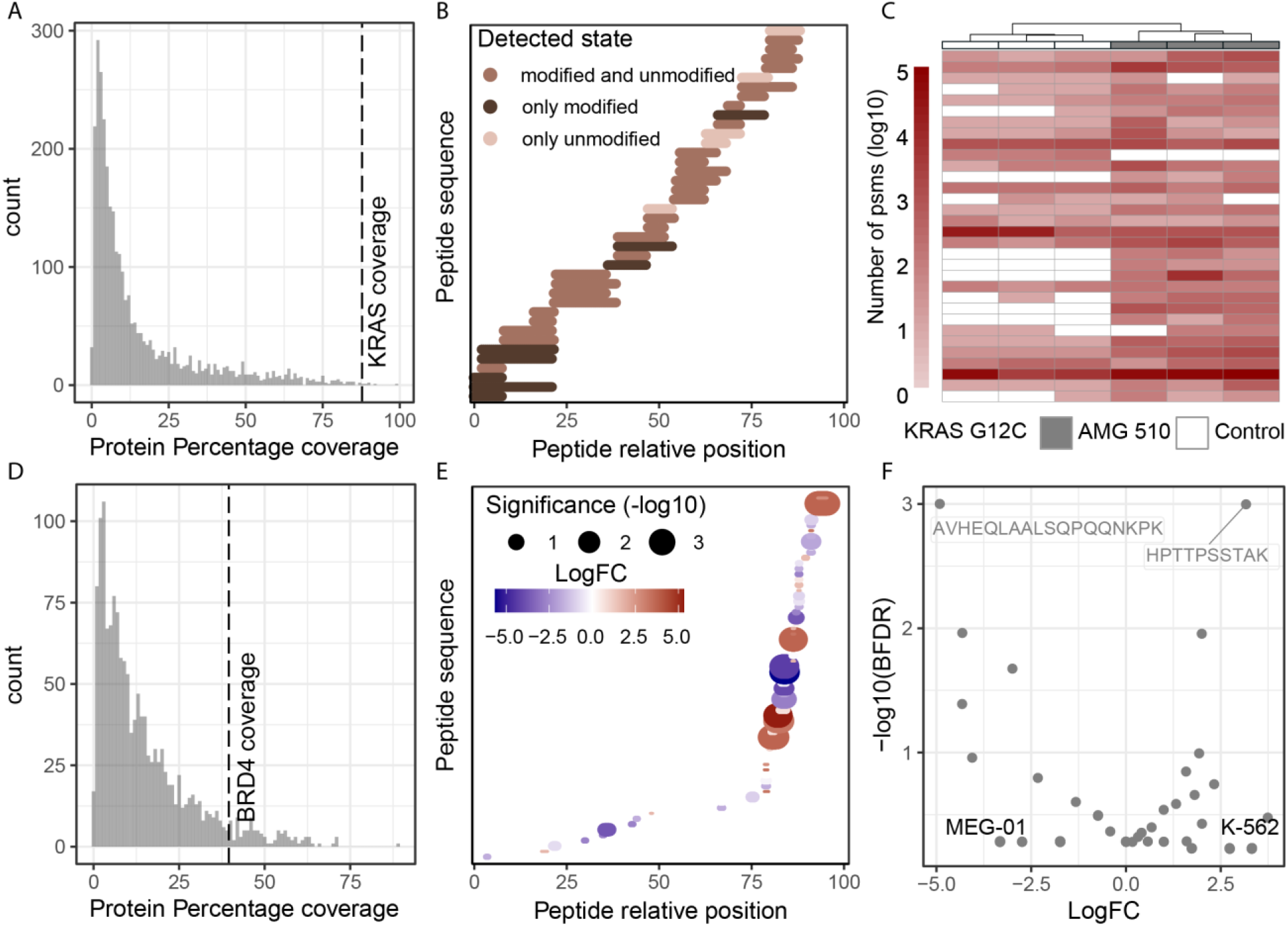
a. Percentage protein coverage for all preys and baits calculated as the percentage of amino acids detected per gene according to the longest isoform, across all KRAS IP-MS samples and all peptidoforms. Dotted line highlight the bait (KRAS) protein coverage. b. Relative position of all detected KRAS bait peptides, as well as whether they were detected in unmodified, modified or in multiple delta masses. Peptides with isotopic error delta masses were considered unmodified for this representation. c. Clustering of post-normalised PSM counts for highly abundant peptidoforms belonging to bait KRAS G12C protein. Unsupervised clustering performed with default hclust parameters with peptidoforms not detected in specific samples assigned a zero value following log transformation. d. Percentage protein coverage for all preys and baits calculated as the percentage of amino acids detected per gene according to the longest isoform, across all BRD4 IP-MS samples and all peptidoforms. Dotted line highlight the bait (BRD4) protein coverage e. Relative positions of BRD4 bait peptides between K-562 and MOLM-13, with peptides with statistical significance and fold change encoded in size and color respectively. f. Differential bait BRD4 peptidoforms detected between K-562 and MEG-01 samples according to the SAINTexpress, defined as BFDR< 0.05. Only bait peptidoforms were included in the SAINTexpress analysis.

## References

1. Smith, L. M. et al. Proteoform: a single term describing protein complexity. Nat Methods 10, 186 (2013).

2. Melani, R. D. et al. The Blood Proteoform Atlas: A reference map of proteoforms in human hematopoietic cells. Science (1979) 375, 411–418 (2022).

3. Toby, T. K. et al. Proteoforms in Peripheral Blood Mononuclear Cells as Novel Rejection Biomarkers in Liver Transplant Recipients. Am J Transplant 17, 2458 (2017).

4. Demeulemeester, N. et al. msqrob2PTM: differential abundance and differential usage analysis of MS-based proteomics data at the post-translational modification and peptidoform level. Molecular & Cellular Proteomics 100708 (2023) doi:10.1016/J.MCPRO.2023.100708.

5. Trinkle-Mulcahy, L. et al. Identifying specific protein interaction partners using quantitative mass spectrometry and bead proteomes. J Cell Biol 183, 223 (2008).

6. Yu, F. et al. Identification of modified peptides using localization-aware open search. Nature Communications 2020 11:1 11, 1–9 (2020).

7. Kong, A. T., Leprevost, F. V., Avtonomov, D. M., Mellacheruvu, D. & Nesvizhskii, A. I. MSFragger: ultrafast and comprehensive peptide identification in shotgun proteomics. Nat Methods 14, 513 (2017).

8. Teo, G. et al. SAINTexpress: improvements and additional features in Significance Analysis of Interactome software. J Proteomics 100, 37 (2014).

9. Swaney, D. L. et al. A protein network map of head and neck cancer reveals PIK3CA mutant drug sensitivity. Science 374, eabf2911 (2021).

10. Lanman, B. A. et al. Discovery of a Covalent Inhibitor of KRASG12C (AMG 510) for the Treatment of Solid Tumors. J Med Chem 63, 52–65 (2020).

11. Salmón, M. et al. Kras oncogene ablation prevents resistance in advanced lung adenocarcinomas. J Clin Invest 133, (2023).

12. Nolan, A. et al. Proteomic Mapping of the Interactome of KRAS Mutants Identifies New Features of RAS Signalling Networks and the Mechanism of Action of Sotorasib. Cancers (Basel) 15, 4141–4141 (2023).

13. Huang, L., Carney, J., Cardona, D. M. & Counter, C. M. Decreased tumorigenesis in mice with a Kras point mutation at C118. Nature Communications 2014 5:1 5, 1–10 (2014).

14. Kramer-Drauberg, M. & Ambrogio, C. Discoveries in the redox regulation of KRAS. Int J Biochem Cell Biol 131, 105901 (2021).

15. Boja, E. S. & Fales, H. M. Overalkylation of a protein digest with iodoacetamide. Anal Chem 73, 3576–3582 (2001).

16. Zhang, K., Li, Y., Huang, T. & Li, Z. Potential application of TurboID-based proximity labeling in studying the protein interaction network in plant response to abiotic stress. Front Plant Sci 13, 974598 (2022).

17. Sharifi Tabar, M. et al. Illuminating the dark protein-protein interactome. Cell Reports Methods 2, 100275 (2022).

18. Sdelci, S. et al. MTHFD1 interaction with BRD4 links folate metabolism to transcriptional regulation. Nat Genet 51, 990 (2019).

19. Roux, K. J., Kim, D. I., Raida, M. & Burke, B. A promiscuous biotin ligase fusion protein identifies proximal and interacting proteins in mammalian cells. Journal of Cell Biology 196, 801–810 (2012).

20. Van Andel, E. et al. Highly Specific Protein Identification by Immunoprecipitation-Mass Spectrometry Using Antifouling Microbeads. ACS Appl Mater Interfaces 14, 23102–23116 (2022).

21. Perez-Riverol, Y. et al. The PRIDE database resources in 2022: a hub for mass spectrometry-based proteomics evidences. Nucleic Acids Res 50, D543–D552 (2022).

22. Geiszler, D. J. et al. PTM-Shepherd: Analysis and Summarization of Post-Translational and Chemical Modifications From Open Search Results. Mol Cell Proteomics 20, 100018 (2021).

23. Land, H. & Humble, M. S. YASARA: A Tool to Obtain Structural Guidance in Biocatalytic Investigations. Methods Mol Biol 1685, 43–67 (2018).

24. Berman, H. M. et al. The Protein Data Bank. Nucleic Acids Res 28, 235–242 (2000).

25. Delgado, J., Radusky, L. G., Cianferoni, D. & Serrano, L. FoldX 5.0: working with RNA, small molecules and a new graphical interface. Bioinformatics 35, 4168–4169 (2019).

